# The Effect of Population Structure on Murine Genome-Wide Association Studies

**DOI:** 10.1101/2020.09.01.278762

**Authors:** Meiyue Wang, Zhuoqing Fang, Boyoung Yoo, Gill Bejerano, Gary Peltz

## Abstract

Population structure (PS) has been shown to cause false positive signals in genome-wide association studies (GWAS). Since PS correction is routinely used in human GWAS, it was assumed that it also should be utilized for murine GWAS using inbred strains. Nevertheless, there are fundamental differences between murine and human GWAS, and the impact of PS on murine GWAS results has not been thoroughly investigated. To assess the impact of PS on murine GWAS, we examined 8223 datasets that characterized biomedical responses in panels of inbred mouse strains. Surprisingly, we found that the PS had a minimal impact on datasets characterizing responses in ≤20 strains; and relatively little impact on the majority of datasets characterizing >20 strains. Moreover, there were examples where association signals within known causative genes could be rejected if PS correction methods were utilized. PS assessment should be carefully used and should be considered in conjunction with other criteria when assessing the candidate genes identified in GWAS using inbred mouse strains.

## Introduction

Because of ancestral relatedness among the individuals within an analyzed population, a GWAS will identify a true causative genetic variant along with multiple other false positive associations, which arise because of genetic regions that were commonly inherited within a sub-population. This property, which is referred to as ‘population structure’ (PS) has been shown to exist in populations ranging from plants [1] to humans [2, 3]; and it inflates the number of false positive results obtained in a GWAS. Since PS was identified as a significant confounding factor for human GWAS, methods were developed to distinguish the false positive PS-based associations from the true causative genetic factors for a studied trait. A unified mixed model method was developed [2] to control for PS using a matrix [4] where Bayesian clustering is used to infer the number of subpopulations and to estimate the effect of population structure. Improved methods were subsequently developed over the next decade. The matrix was replaced by the use of principal components (PC) that summarized the genome-wide patterns of relatedness [1], and principal component analysis (PCA) was shown to be useful for inferring PS from genetic data [5, 6]. PCA has two advantages over the population structure matrix: (i) the subpopulations do not have to be specified prior to the analysis, which can be an arbitrary process that introduces errors; and (ii) it is far more computationally efficient, which is important when a large number of individuals with many SNPs are evaluated.

Although the methodology used to assess PS has improved, we do not know whether PS has a significant impact on GWAS results using inbred mouse strains. Mouse is the premier model organism for biomedical discovery, and many therapies were initially discovered using mice. Since the inbred laboratory strains are derived from what is estimated to be four ancestral founders that diverged ~1 million years ago [7, 8], PS could certainly impact murine GWAS results. Others have advocated that PS correction should be used in murine GWAS [9, 10]. However, murine and human GWAS differ in several fundamental ways. A typical human GWAS includes thousands of individuals collected from a natural population. In contrast, most murine GWAS use less than 50 inbred strains of known ancestry [11], the strains are homozygous, they do not interbreed, and environmental and other variables are tightly controlled. Because of this, genetic effect sizes in murine GWAS are much larger than in human GWAS. Because of these differences, we examined a large database of responses measured in panels of inbred strains to assess the impact of PS on GWAS outcome. For this analysis, we analyzed results obtained using haplotype-based computational genetic mapping (HBCGM) - a method for performing GWAS in mice - that has important differences from conventional SNP-based studies [12]. In an HBCGM experiment, a property of interest is measured in available inbred mouse strains whose genomes have been sequenced; and genetic factors are computationally predicted by identifying genomic regions (haplotype blocks) where the pattern of within-block genetic variation correlates with the distribution of phenotypic responses among the strains [12–14]. HBCGM analyses use a SNP database with 25M SNPs with alleles covering 49 inbred mouse strains (**Table S1**) generated from analysis of whole genome sequence data [15].

HBCGM has successfully identified genetic factors for >22 biomedical traits in mice [12, 13, 16–37]. However, as with other GWAS methods, HBCGM analyses identify many genomic regions with allelic patterns that correlate with a phenotypic response pattern; but only one (or a few) contains a causative genetic factor [12]. Therefore, we investigated the effect of PS on murine GWAS results, and the utility of applying a PS association test for eliminating false positives from the list of candidate genes identified by HBCGM.

## Results

The Mouse Phenome Database (**MPD**) (https://phenome.jax.org) [38] contains 8223 datasets that characterize basal, age-related, and experimentally-induced responses (i.e. ‘phenotypes’) in panels of inbred mouse strains. This database has a total of 1.52M individually measured responses. We previously demonstrated that MPD datasets have utility for genetic discovery; a genetic susceptibility factor for a drug-induced CNS toxicity was identified by HBCGM analysis of one MPD dataset [21]. Therefore, we initially selected all MPD datasets that measured a response in 10 or more strains whose genomic sequence was available (2435 datasets). For each of these datasets, haplotype blocks with allelic patterns that correlated with the measured strain response pattern were identified by HBCGM. The average number of correlated blocks (*p_HBCGM_* < 0.01) for each dataset was 3966, which were selected from among the 6 to 50 million haplotype blocks produced by the algorithm for each dataset. The number of blocks assembled by the algorithm depended upon the number of strains contained in a dataset. We then wanted to use a multi-variate association test (MANOVA) to determine whether the haplotypic strain groupings within the correlated blocks were related to PS among the analyzed strains. However, to do this, the number of PCs has to be specified in advance to perform the PS association test. Therefore, we first examined the percentage of the variance that was explained when a variable number of principal components (PCs), which ranged from 1 to 33 because <33 inbred strains were analyzed in any dataset, were used for the PCA analysis. Because the curves on the Scree plots for most of the 2435 datasets evaluated had a bend (i.e. ‘elbow’) that occurred between the 3^rd^ and 5^th^ PC, we used the first four PCs (total genetic variance ranged between 26%-59%) as the response variable that was used for the PS association analyses (**Fig. S1**).

A pairwise identity-by-state (IBS) matrix divided the 49 sequenced inbred strains into four sub-populations (**Table 1, Fig. 1**). These strain groupings are based upon their genome wide genetic relatedness, and their sub-population grouping provided a pre-determined label that is used in the subsequent analyses. Sub-populations 2 and 3 contain the majority of the inbred strains, which are closely related. The sub-population 1 strains are derived from a C57BL ancestor; and the five strains in sub-population 4 are genetically distinct from the other groups. The spatial relationship of the 49 strains (plotted using the first two PCs for each strain) is concordant with the hierarchical clustering (**Fig 1)**. A separately performed quantitative analysis, which used the EIGENSOFT/smartpca program [39] to generate Tracy-Widom (TW) statistics and ANOVA values for the groupings, confirms that two PCs capture the PS for these strains (**Tables S2A, 2B**).

**Figure 1.**
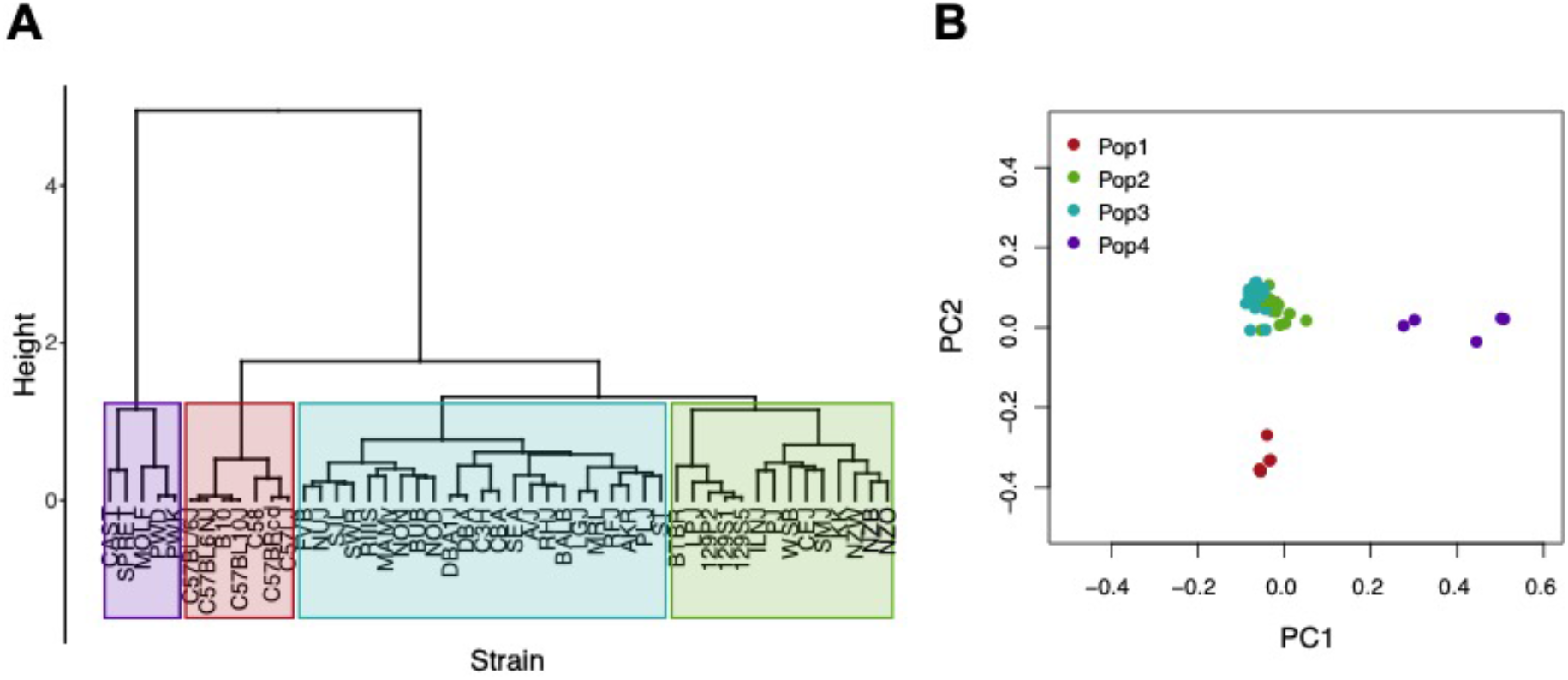
An analysis of population structure among 49 inbred mouse strains, which is based upon whole genome sequence analysis, identifies four sub-populations. (**A**) The relatedness of the 49 inbred strains based upon hierarchical clustering using an identity-by-state similarity matrix; or (**B**) a scatter plot generated by PCA using the first two PCs for each strain are shown. The sub-populations identified by the two methods are completely concordant. Sub-populations 1 and 4 are distinct from the majority of the inbred strains that contained in two closely related sub-populations (2 and 3).

**Table 1.** The 49 inbred strains can be divided into the four groups shown in this table based on their pattern of genome-wide allelic sharing.

| Group | Number of Strains | Strain List |
| --- | --- | --- |
| 1 | 7 | C57BL/6J, B10, C57BL10J, C57BL6NJ, C57BRcd, C57LJ, C58 |
| 2 | 14 | BTBR, CEJ, KK, NZB, NZW, 129P2, 129S1, 129S5, ILNJ, LPJ, NZO, PJ, SMJ, WSB |
| 3 | 23 | BUB, DBA1J, FVB, NON, NUJ, RFJ, RHJ, RIIIS, SJL, A/J, AKR, BALB, C3H, CBA, DBA, LGJ, MAMy, MRL, NOD, PLJ, SEA, ST, SWR |
| 4 | 5 | CAST, MOLF, PWD, PWK, SPRET |

We then examined PS among the strain panels used in the MPD datasets. The number of inbred strains analyzed in each of the 2435 MPD datasets with data for >10 evaluable strains are summarized in **Table S3**. During our analysis, we noted that many different MPD datasets used the same panel of inbred strains, which is because multiple phenotypes were evaluated by the same investigator, and because certain strains are commonly used by different laboratories. Therefore, we could examine PS among the strains used in the majority (55%) of the 2345 MPD datasets by examining the 22 sets of inbred strains that were repeatedly used in many MPD datasets (**Table S2C**). Our initial analysis of the PS graphs generated using two PCs indicated that we should not assess population structure in MPD datasets that analyzed < 20 strains because: (i) the population substructure was extremely variable, and (ii) the strain groupings within these datasets often contained strains from different global sub-groups (**Fig. S2**). To confirm the visual observations, we also used the EIGENSOFT/smartpca program to analyze the PS in the panels with < 20 inbred strains, since it provides an unsupervised analysis that ignores the pre-determined of sub-population for each strain. The results indicated that the strain groupings did not have significant PS: all Tracy-Widom (TW) p-values were far above 0.05 for the first two PCs (**Table S2C**). It is also noteworthy that the TW p-values decreased as the strain number increased, which indicates that it is easier to identify PS among groups with a larger number of strains. Overall, only 3 of the 22 strain panels evaluated had a TW p-value <0.05 for the first PC, and all 3 of those panels had >29 inbred strains (Table S2C). These results indicated that most strain panels used in the MPD either do not have PS that needs to be corrected or the PS among the strains is not large enough for PCA to capture it.

We further examined population sub-structure in 1750 MPD datasets that examined responses in > 20 inbred strains. To illustrate the general properties that emerged from our analyses, we show 960 MPD datasets that repeatedly analyzed responses in the same sets of (n=23-32) inbred strains. The first two PCs for 432 of these datasets did not identify significant PS; there were no clear groupings for the strains; and the TW p-values are all >0.05, which may be due to the small number of strains analyzed (**Fig. S3, Table S2C**). In contrast, the PCA plots indicated that PS could be present in 528 other MPD datasets (**Fig. S4**) where the group 1 strains (C57BL related) are clearly separated from the other strains. However, in these datasets, the global group 2 and group 3 strains are broadly distributed in the graphs, without an explicit boundary that separates them from the other strains; and the groups 2 and 3 strains are intermixed with group 1 strains in many of these graphs. It should be noted that 256 of these 528 datasets use two recurring strain panels: 178 datasets use same 24 strain panel and 78 datasets use the same 25 strain panel (**Fig. S4A-B**). For both of these recurring panels, the TW p-values indicate clearly what they show (**Table S2C**). Of importance, even when datasets examine responses in strain panel that appear to have PS, it will only have an effect if the phenotypic response pattern completely mirrors that of the global strain sub-populations. If this type of response pattern does not occur, which appears to be the case for the majority of the measured responses (see below), PS would have a limited effect on genetic analysis results.

To more directly assess PS impact on the haplotype blocks generated by HBCGM analysis of the 2435 MPD datasets with >10 strains, a PS association test was performed on each correlated haplotype block. An adjusted p-value for the PS association test for each block was generated using MANOVA. Blocks with a *p_adj_* ≤ 0.05 have a significant association with population structure (i.e. PS^+^), and could be removed from further consideration, while those with a *p_adj_* ≥ 0.05 are viewed as viable candidate genes for further evaluation (PS^-^). For 68% of the datasets (1,660 of 2435 analyzed), >50% of the correlated blocks were not associated with population structure (PS^-^); and 39% of the datasets (949 of 2435) had 75 to 100% PS^-^ blocks (**Fig. 2**). Only 32% of the datasets (n=775) had >50% PS^+^ correlated blocks; and most of these (23%, 565 datasets) have between 25 and 49% PS^-^ blocks. Only 9% of the MPD datasets (n= 210) have >75% PS^+^ blocks. Overall, our results indicate that for most MPD datasets, the vast majority of the haplotype blocks identified by HBCGM are not affected by PS. We also investigated whether the magnitude of the PS impact is affected by the number of strains analyzed (i.e. the sample size). As the strain number increased, the number of correlated candidate blocks identified by HBCGM analysis increased (**Fig. 3A**). This result is consistent with prior studies indicating that genetic analyses, which are performed on large populations, will identify additional genetic variants with a small effect size [40]. However, while the number of PS^-^ blocks plateaued after 15 strains were analyzed, the number of PS^+^ blocks increased as the number of analyzed strains increased (**Figs. 3B-C**). These results indicate that when an increased number of inbred strains are analyzed, the number of correlated haplotype blocks increases, as does percentage of PS^+^ blocks. The results are completely consistent with the sample size effects previously noted in human-case control studies [3].

**Figure 2.**
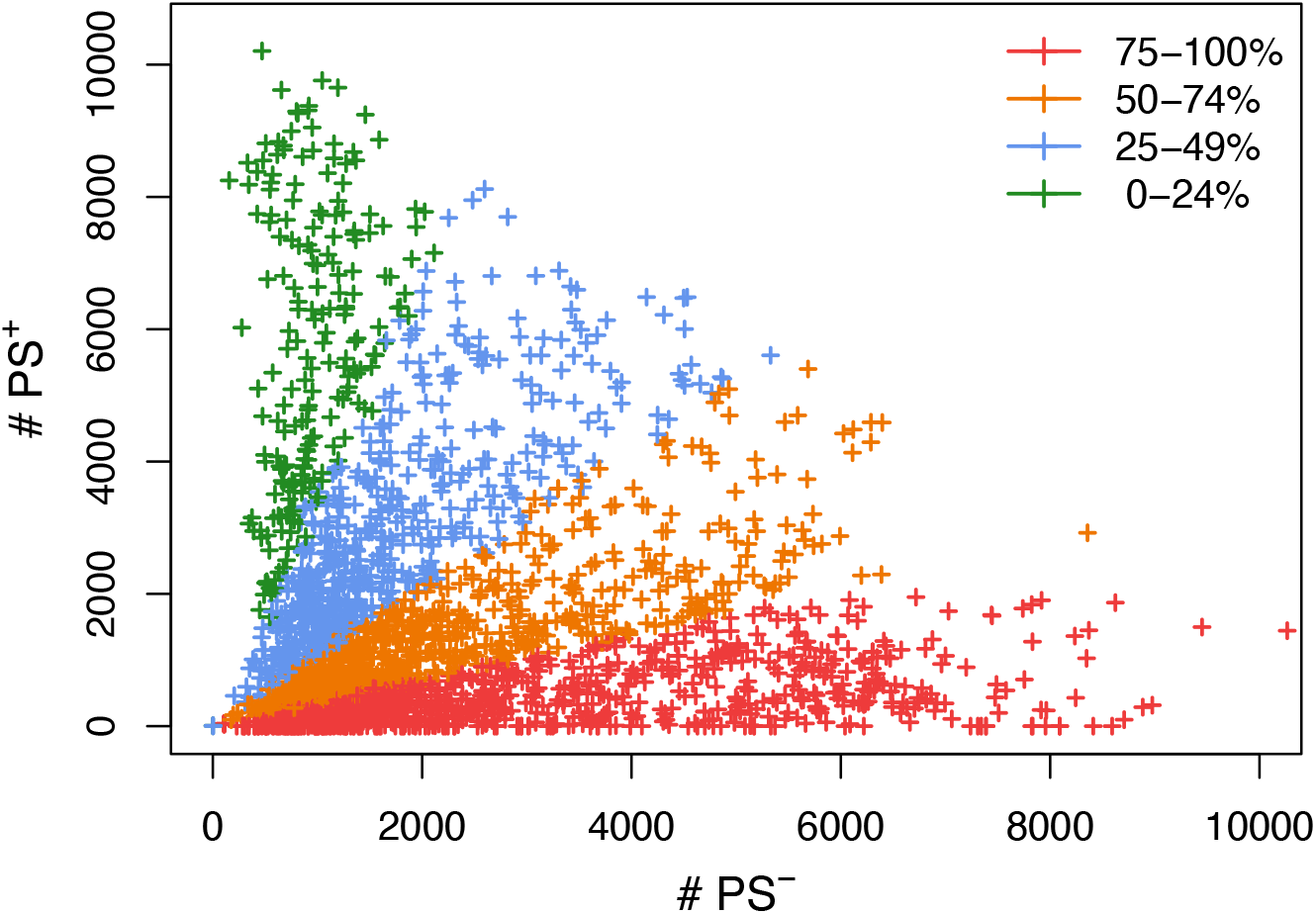
A scatter plot showing the number of candidate haplotype blocks associated with population structure (PS^+^) relative to PS^-^ candidate blocks. After 2435 MPD datasets were analyzed by HBCGM, candidate blocks (*p_HBCGM_* < 0.01) were analyzed by an association test to determine whether they were related to population structure among the inbred strains that were analyzed. Each datapoint (+) indicates the number of PS^+^ (y-axis) and PS^-^ (x-axis) blocks identified for one MPD dataset. There are 949 MPD datasets where 75% to 100% of the blocks are PS^-^ (shown in red); the 711 datasets with 51-74% PS^-^ blocks are shown in orange; and the 565 datasets with 25-49% PS^-^ haplotype blocks are shown in blue; and the 210 datasets with 0-24% PS^-^ blocks are shown in green.

**Figure 3.**
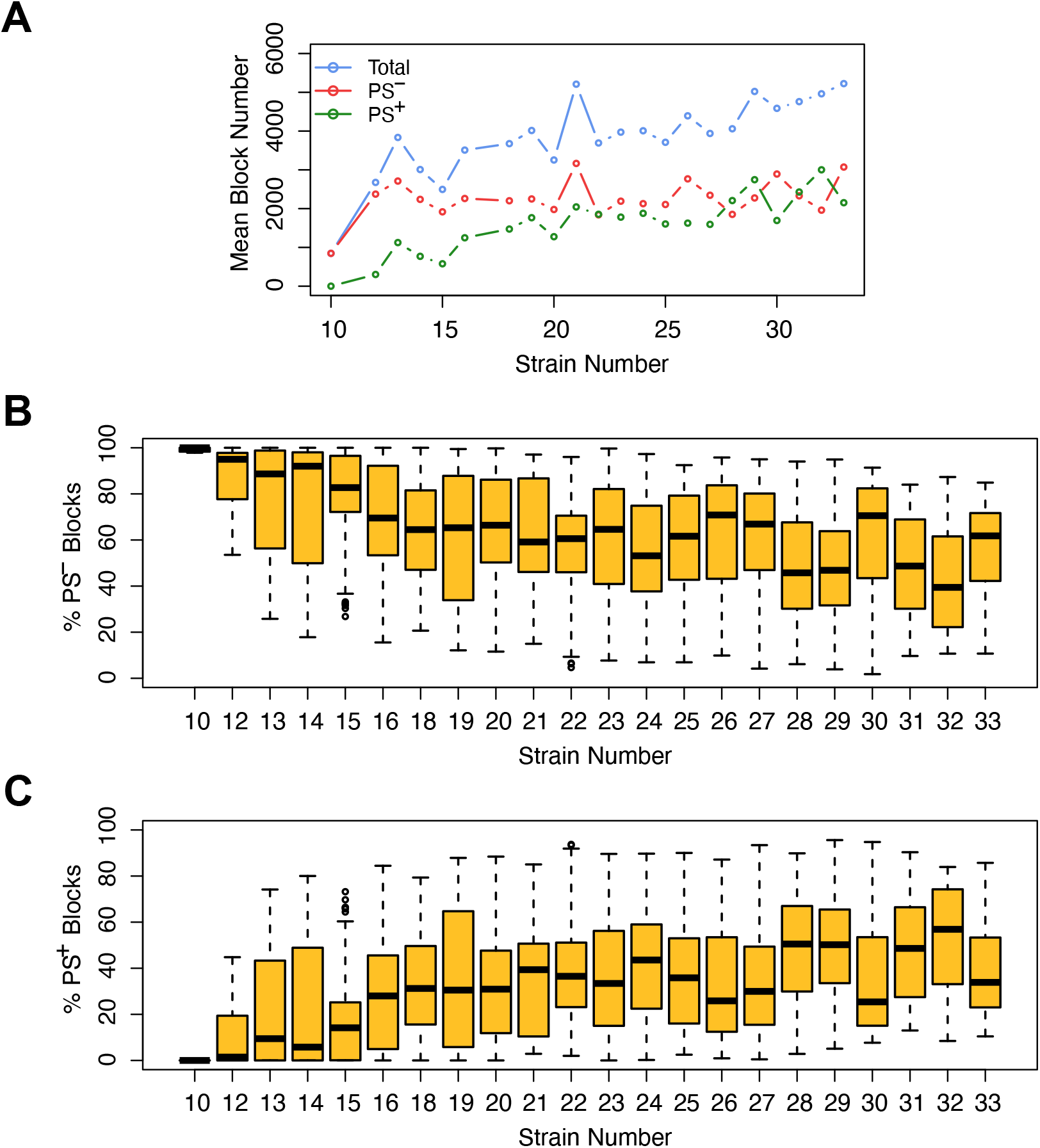
The effect of population structure increases with the number of analyzed strains. Analysis of the total number of candidate haplotype blocks, the number of blocks with population structure (PS^+^), and the number of PS-independent (PS^-^) blocks are shown as a function of the number of analyzed strains. After 2435 MPD datasets were analyzed by HBCGM, the correlated blocks (*p_HBCGM_* < 0.01) were analyzed by an association test to determine whether population structure had a significant influence on the strain groupings within the blocks. (**A**) The results were then graphed as a function of the number of mouse strains within each dataset (range 10 – 33). A blue circle represents the average of the total number of candidate blocks, and the mean number of PS^-^ (red) and PS^+^ blocks (green) are also shown in this graph. (**B, C**) The percentage of (**B**) PS^-^ and (**C**) PS^+^ blocks were then assessed for each dataset. The box plots indicate the 25th and 75^th^ percentile, and the black bar indicates the median value. While the number of PS^-^ blocks plateaued after 15 strains were analyzed, the number of PS^+^ blocks increased in the datasets that analyzed an increased number of strains.

When considering whether PS correction should be utilized for assessing mouse GWAS results, it is important to determine whether this could lead to rejection of a true causative association. Therefore, we investigated whether PS was present in haplotype blocks within genes whose allelic patterns are known to be causal for phenotypic response differences in 6 MPD datasets (**Table 2**). We first examined the haplotype blocks identified by HBCGM from analysis of data on strain susceptibility to anthrax toxin (MPD 1501), which is known to be caused by allelic variation within the *Nalp1a* and *Nalp1b* genes [41]. Both of the correlated haplotype blocks within these genes were PS^-^. Similarly, the identified haplotype block within an experimentally validated causative gene (*Abcb5*) [21] affecting susceptibility to a drug (haloperidol)-induced CNS toxicity (MPD 39410) also was PS^-^. The albino skin type (MPD 22001) that appears in some inbred strains is determined by a *Cys103Ser* SNP within the tyrosinase (*Tyr*) gene [42], and the correlated haplotype block identified by HBCGM analysis within *Tyr* was also PS^-^.

**Table 2.** The results of PS analysis performed on haplotype blocks within known causative genes for 6 MPD datasets are shown. The MPD dataset number, the sex of the mice, a description of the measured response, and the number of strains analyzed in that dataset are shown. The gene symbol for the causative gene, the chromosome and position of the identified haplotype block, and the p-value and adjusted p-value for the PS association test for that block are shown.

| MPD Dataset | Number of Strains | Gene | Block position | HBCGM p-val | PS p-val | PS adj p-val |
| --- | --- | --- | --- | --- | --- | --- |
| 1501 F susceptibility to *Bacillus anthracis* | 23 | *Nlrp1a* | Chr11: 71110959-71111942 | 0 | 0.9537 | 0.9702 |
| 1501 F susceptibility to *Bacillus anthracis* | 23 | *Nlrp1b* | Chr11: 71159763-71159873 | 0 | 0.5785 | 0.6782 |
| 26721 F retinal degeneration | 29 | *Pde6b* | Chr5: 108399551-108400383 | 0 | 0.0244 | 0.0491 |
| 26721 M retinal degeneration | 29 | *Pde6b* | Chr5: 108399551-108400383 | 0 | 0.0244 | 0.0491 |
| 9904 F HDL cholesterol baseline | 30 | *Apoa2* | Chr1: 171225795-171225890 | 5.5e-6 | 0.0005 | 0.0010 |
| 9904 M HDL cholesterol baseline | 31 | *Apoa2* | Chr1: 171225644-171225697 | 3.14e-5 | 0.1537 | 0.2448 |
| 9907 F HDL cholesterol after 17 wks on diet | 30 | *Apoa2* | Chr1: 171227457-171227593 | 0.0066 | 0.0039 | 0.0106 |
| 9907 M HDL cholesterol after 17 wks on diet | 25 | *Apoa2* | Chr1: 171227457-171227593 | 0.0008 | 0.0020 | 0.0044 |
| 22001 F coat color | 42 | *Tyr* | Chr7: 87446687-87446831 | 3.68e-6 | 0.1071 | 1 |
| 22001 M coat color | 42 | *Tyr* | Chr7: 87446687-87446831 | 3.68e-6 | 0.1071 | 1 |
| 39410 M Haloperidol induced latency day 30 | 18 | *Abcb5* | Chr12: 118885164-118916966 | 4.19e-10 | 0.1507 | 0.8047 |

However, the results of PS analyses for three other MPD datasets raised concerns. *Apoa2* encodes the second most abundant protein within high density lipoprotein (HDL) particles, and it is involved in lipoprotein metabolism. *Apoa2* alleles were previously associated with differences in plasma HDL cholesterol levels in mice [43]; and HDL levels were 70% decreased in *Apoa2* knockout mice [44]. HBCGM analysis of two datasets measuring HDL cholesterol levels (MPD 9904 and 9907) indicated that 3 of 4 correlated haplotype blocks within *Apoa2* are PS^+^ blocks (MANOVA *p_adj_* < 0.05). Retinal degeneration in inbred strains is known to be caused by a stop codon (*Tyr347X*) within *phosphodiesterase 6b (Pde6b*) [45]. One MPD dataset (MPD 26721) examined the retinas of 29 inbred strains: 21 strains had normal retinas, and 8 strains had retinal degeneration. HBCGM analysis identified two *Pde6b* haplotype blocks that completely correlated with retinal degeneration in male and female mice (*p_HBCGM_* = 0). However, the strain groupings within these blocks had PS; the PS association test p-values for these blocks were 0.02 (*p_adj_* = 0.049) (Table 2). The blocks had PS because all 8 strains with retinal degeneration were from population group 3, and all population group 1 and 2 strains had normal retinas. However, several group 3 strains had normal retinas and *Pde6b Try347* alleles (**Fig. 4**). These examples demonstrate that some true positives, if the usual FDR control rate (*q* = 0.05) was applied, could have been falsely rejected based upon their association with PS.

**Figure 4.**
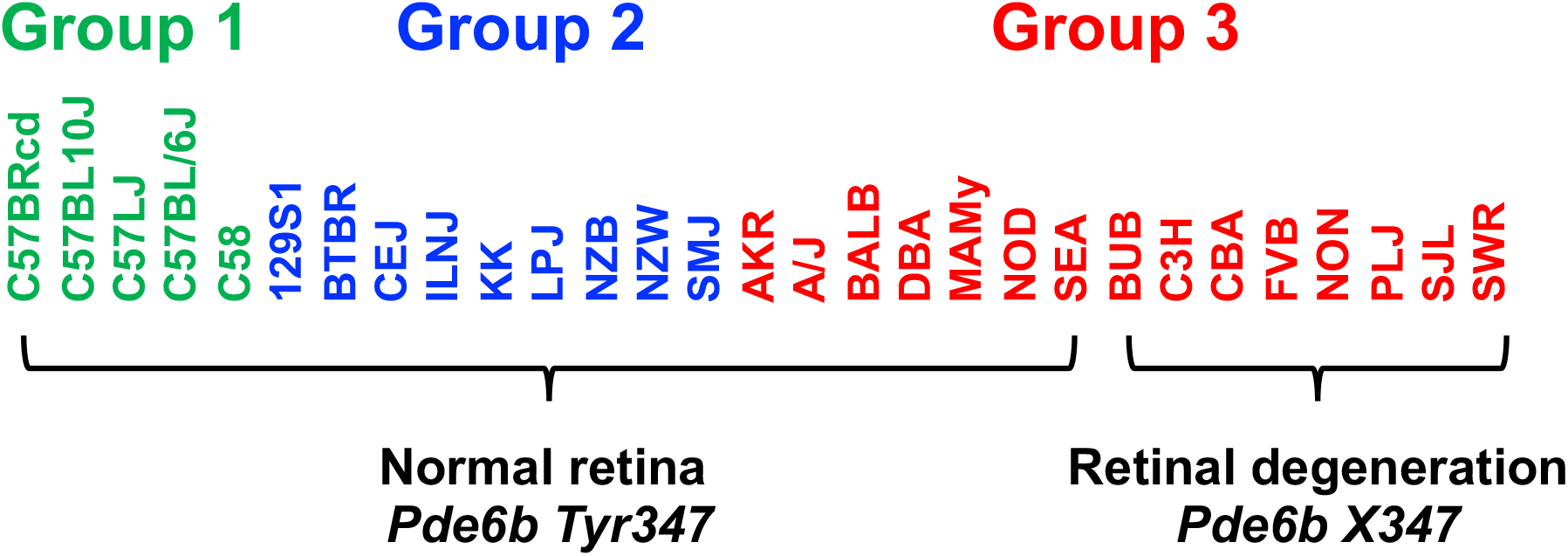
The haplotype block with a causative mutation is associated with population structure. MPD 26721 examined the retinas of 29 inbred strains: 21 strains had normal retinas and 8 strains had retinal degeneration. A haplotype block within *Pde6b* contained the causative SNP (*Tyr347X*) for this type of retinal degeneration. All strains with retinal degeneration had the *Pde6b 347X* allele, while those with normal retinas had the *Tyr347* allele. The haplotype block had PS, because all group 1 and 2 strains (based upon hierarchical clustering of whole genome sequence data from 49 inbred strains (**Table 1**)) had normal retinas; while all strains with retinal degeneration were group 3 strains. However, several group 3 strains (AKR, A/J, BALB, DBA, MaMy, NOD, SEA) had normal retinas and the *Tyr347* allele. Thus, while the strain groupings within the block have PS based upon their global allele sharing pattern, the allelic pattern within the haplotype block had a stronger association with retinal degeneration.

## Discussion

PS correction is commonly performed when analyzing GWAS results involving human or other species (cattle, maize, etc.), and PS correction has also been advocated for use in murine GWAS [9, 10]. While PS correction helps to eliminate false positives, our analyses indicate that PS makes a smaller than expected contribution to most murine GWAS studies. Moreover, we found that PS correction can even generate a false negative result, i.e. it can lead to rejection of an experimentally confirmed true causative genetic factor. Of importance, this analysis evaluated the largest available dataset of phenotypic information for inbred mouse strains, and it was assembled using data that was obtained from the vast majority of investigators studying genetic traits in mice. *Why is the utility of PS correction in murine HBCGM analyses different from that for association studies performed using different methods or involving other species?* We identify four factors that account for this difference. (i) A very limited number of inbred strains are examined in a murine GWAS, which usually analyze <20 (and never >40 inbred strains). This is orders of magnitude less than the number of subjects in human GWAS (now ranging from thousands to hundreds of thousands). Since the PS effect increases as the number of inbred strains analyzed are increased, PS has a more limited effect on most murine GWAS. (ii) We found that the vast majority of murine GWAS studies utilize strains with limited PS. Most (37 of 49 or 75%) of the commonly used inbred strains are derived from closely related populations, which have limited or no population structure. Among 25M SNPs that were analyzed, pairwise comparisons revealed that the level of allelic similarity among the classical inbred strains is >70%. The limited amount of genetic variation among these strains precludes their separation into distinct sub-populations. (iii) Human (and other species) GWAS identify trait associations using SNP markers, and the association signals depend upon the existence of linkage disequilibrium (LD) between SNP markers and causal genetic variants. The dependence upon LD, which extends over a region of indeterminate size, increases the effect that regional PS could have on an outcome. In contrast, HBCGM does not rely on LD between marker and causative SNPs. HBCGM uses a combination of adjacent SNPs to produce haplotype blocks, which are the composite genetic variants that are analyzed. Since haplotype blocks are assembled from analysis of whole genome sequence, the block boundaries are precisely determined, and the analyzed variants contain the causative genetic factors. (iv) The impact of a false negative result (excluding a true positive due to PS) is much greater for a murine GWAS. Genetic association studies involving large populations usually identify many genetic variants, with each having a small genetic effect size. In those situations, the loss of a few true positive associations does not create a large problem since many others remain. However, murine GWAS always analyze a small number of inbred strains; and the heritability and genetic effect size of the identified candidate genes is relatively large (usually >0.3) because their genome is homozygous and environmental and other confounding factors are minimized. Thus, unlike its small effect on human GWAS results, the elimination of a true positive by PS correction can be disastrous for a murine GWAS. Of note, only one of the above factors is specific to HBCGM, while the other three factors are relevant to all forms of murine GWAS.

We have shown in several situations that a true causative factor could be associated with strain phylogenetic background. In two examples, *Apoa2* (MPD 9904) and *Pde6b* (MPD 1501), PS correction could have removed true causative blocks from further consideration. However, in one case (retinal degeneration and *Pde6b),* the identified haplotype block was much more strongly associated with the phenotypic response pattern (genetic association p-value=0) than with population sub-structure (p-value=0.49). In another case (HDL levels and *Apoa2),* the p-values for the genetic and the population structure association tests for the causative haplotype block were of a similar magnitude, but published information indicated that the gene candidate was very strongly associated with the analyzed phenotype. As suggested by others after examining GWAS results for multiple traits in plants, it is not easy to distinguish between a true and a spurious association due to genetic background, even after correcting for PS [45]. However, when GWAS are performed under conditions with true genome wide coverage, a true association is expected to exhibit the strongest association [46]. Allele sharing within a localized candidate genomic region should be greater than one based upon genome wide allelic correlations. Thus, examining the ratio of the p-values obtained from the GWAS and PS association tests could provide a more informative way to eliminate spurious positives while retaining the true positive associations. Nevertheless, as was previously observed in plants [46], there are situations (as with HDL and *Apoa2*) where a shared strain background can be responsible for trait response differences. In these situations, the strength of the functional evidence that a candidate gene could be responsible for a trait difference could override PS considerations. Various recombinant inbred (RI) strain panels have been used for genetic mapping studies in mice, which include: the Hybrid Mouse Diversity Panel [47], which has 100 RI strains generated from 30 founder strains; the Diversity Outbred [48] and Collaborative Cross [49] panels were generated from 8 founder strains (A/J, C57BL/6J, 129S1/SvImJ, NOD/ShiLtJ, NZO/HlLtJ, CAST/EiJ, PWK/PhJ, and WSB/EiJ); and the BXD RI panel [50], which was generated from 2 founder strains (C57BL/6 and DBA/2). Of importance, all of the founder strains that were used to generate these RI panels are a subset of the strains evaluated here. Thus, our cautions about the utilization of PS correction methods may be relevant to studies performed using these RI panels.

Lastly, other methods can be used to eliminate false positive associations in GWAS. We have shown that true positives can be identified by the use of orthogonal criteria for analyzing HBCGM output. Causative genetic factors were selected from among the many genes with correlated genetic patterns using gene expression and metabolomic data [28], curated biologic information [51], or the genomic regions delimited by prior QTL analyses [30, 31]. This integrated approach evaluates genetic candidates using multiple criteria, and it can produce results that are superior to that of using a single highly stringent genetic criterion to identify gene candidates. Recent efforts to utilize transcriptome wide association studies [52–54], or functional information [52, 54–57] to select causative loci from among the many SNP sites identified in a human GWAS, or to identify SNPs near *a priori* identified gene candidates in plant GWAS [46] resemble our methods for analyzing HBCGM output. In summary, PS assessment may be one factor that should be used along with multiple other factors to assess a candidate gene, which include the relative strength of the GWAS and PS association results, tissue-specific gene expression criteria, and gene-phenotype relationship based upon information contained within the published literature.

## Methods

### Selection of Mouse Phenome Database datasets

MPD datasets (n=8223) were downloaded on March 24, 2020. We analyzed MPD datasets where the mean phenotypic measurement of each strain was obtained from > 5 mice of each strain. An ANOVA test was also performed to determine if the inter-strain variance was significantly greater than intra-strain variances; and a p-value < 1×10^−10^ was used as the cutoff for dataset selection. Datasets with categorical measurements were excluded from bulk analysis of MPD datasets.

### Haplotype block construction and genetic mapping in mice

The sequence of 49 inbred mouse strains were analyzed as previously described [15, 21]. SNPs were dynamically organized into haplotype blocks for each dataset, which only used alleles for the strains contained within the dataset, according to the “maximal” block construction method [24]. In brief, this method produces haplotype blocks with a minimum of 4 SNPs; and each block is only allowed to a predetermined number of haplotypes, which ranges from 2 to 5. Since the “maximal” method enables blocks to overlap, blocks are assembled that cover all possible allelic combinations within a specific genomic region. If a smaller block was nested inside of a larger block and it contained the same haplotypes, it was removed and the larger block was used to cover that region [24]. This ensures that additional SNPs are only included within a block if additional haplotypes are added to the block. The relationship between the phenotypic response pattern and haplotype blocks was evaluated by HBCGM as described [13]. Genes with correlated haplotype blocks were sorted based upon the ANOVA p-value. A cut-off of *p* = 0.01 was used to select haplotype blocks with a correlated allelic pattern. If a gene had multiple correlated blocks, the haplotype block with the smallest p-value was used.

### Population structure association test

We use principal component analysis (PCA) to determine whether a haplotypic strain grouping was associated with PS. Principal components (PC) has been used assess population stratification [6, 39]; it is a major component of the linear mixed model (LMM) that is used to control PS-induced spurious associations in GWAS results. In the LMM, PS is treated as a covariate that influences the phenotypic values in addition to the effect of the genetic markers. However, we treat PS as a dependent variable, which is determined by a comprehensive analysis of genome-wide allelic similarity. For this analysis, the PS of the inbred strains (*y*) is determined by the equation

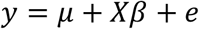

where *y* is an *n* × *p* matrix that is derived from a PCA of sample size of *n* with *p* principal components; *μ* is an *n* × 1 vector that contains the grand mean for each of the *p* variables 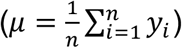; *X* is an *n* × 1 vector of haplotype indicators for *n* strains; *β* is the effect of the haplotype, and *e* is an *n* × 1 vector of the residual error. *p* is a hyperparameter to determine the number of PCs used in analysis, where it guarantees each PC can explain certain amount (say >5%) of the variance of the original genetic relationship. Alternatively, *p* can be arbitrarily selected based upon analysis on a Scree plot (to find the “elbow”), which ranks PCs based on the percentage of variance explained by each PC. If the elbow is observed at *p*-th PC; most of the true signals are captured in the first *p* PCs. By using PC to represent population structure, predetermination of the number of sub-populations is not required.

A multivariate analysis of variance (MANOVA) can be then used to assess the association between strain groupings within a haplotype block and PS, since the strain grouping within a block becomes a single variable that affects the first *p* PCs.

### Generation of genetic relationship and identity-by-state similarity matrices

The genetic relationship matrix (GRM) for inbred mouse strains was generated using genome-wide SNP alleles and GCTA software [58]. The GRM is also known as the variance-covariance standardized relationship matrix, and the eigenvectors of this matrix were used as PC. The GRM eigenvalues for the inbred strains of each PC were used to estimate the amount of GRM variance that PC explains. To assess whether a PC reflects the sub-structure of the GRM, the Tracy-Widom (TW) statistic and corresponding p-values were calculated using EIGENSOFT/smartpca program [39]. This program provides an unsupervised analysis, which ignores the pre-determined global sub-populations identified for each strain. Since we analyze 49 inbred strains whose genomes are homozygous, SNPs were not filtered based upon a minor allele frequency threshold. To further verify that the PCs effectively represent the PS among the strains, we clustered individual strains using a pairwise identity-by-state (IBS) similarity matrix, which was also derived using whole genome SNP data. The IBS similarity matrix is a square, symmetric matrix that reflects the IBS distance between all pairs of inbred mouse strains. PLINK 1.90 [59] was used to calculate the IBS similarity matrix, and it contains values that range from 0 to 1. The hierarchical clustering of 49 strains was determined using the hcut() function within the factoextra/R package (**https://CRAN.R-project.org/package=factoextra**). The sub-population of an inbred strain is based upon its genetic relatedness relative to the other 49 strains. This clustering determines the sub-population for a strain used in subsequent analyses (i.e. their pre-determined label). Then, an ANOVA test is used to evaluate the overall genetic differentiation between any two pre-determined sub-populations along the PCs (i.e. it is a supervised analysis). Hence, the basis for the 4 sub-populations identified using the IBS similarity matrix for the 49 inbred strains can be assessed using the ANOVA test, where the resulting ANOVA p-value is compared with 0.05.

### Multiple test correction for the PS association test

Since the population structure association test was performed on 2435 datasets, the MANOVA test p-value for each block generated by the HBCGM program is adjusted by controlling for the false discovery rate (FDR at *q* = 0.05) using Benjamini-Hochberg method [60]. The adjusted p-value for i-th block is *p_adj_* = *p_i_* × *m/i*, where *p_i_* is the MANOVA test p-value, *m* is the number of blocks (multiple tests), and *i* is the order of *p_i_* in a series of p-values that satisfies *p*_(1)_ ≤ *p*_(2)_ ≤ ⋯ ≤ *p*_(*m*)_. If a block has *p_adj_* ≥ 0.05, it is not considered as having a significant amount of PS (i.e. the null hypothesis, which is that the tested block does not have population structure, cannot be rejected).

## Data availability

The data sets within the Mouse Phenome Database (**MPD**) analyzed in this study are available at (https://phenome.jax.org). All data generated or analyzed during this study are included in this published article [and its supplementary information files].

## Author contributions

The project was formulated at working meetings of all authors. M.W. analyzed the data; and B.Y. and G.B. helped with the analysis. B.Y. and Z.F. contributed code. G.P. and M.W. wrote the paper with input from all authors.

## Abbreviations

GWAS: genome-wide association study
HBCGM: haplotype-based computational genetic mapping
PCA: principal component analysis
PS: population structure

## References

1. Zhao K, Aranzana MJ, Kim S, Lister C, Shindo C, Tang C, et al. An Arabidopsis example of association mapping in structured samples. PLoS Genet. 2007;3(1):e4. Epub 2007/01/24. doi: 10.1371/journal.pgen.0030004. PubMed PMID: 17238287; PubMed Central PMCID: PMCPMC1779303.

2. Yu J, Pressoir G, Briggs WH, Vroh Bi I, Yamasaki M, Doebley JF, et al. A unified mixed-model method for association mapping that accounts for multiple levels of relatedness. Nat Genet. 2006;38(2):203–8. Epub 2005/12/29. doi: 10.1038/ng1702. PubMed PMID: 16380716.

3. Reich DE, Goldstein DB. Detecting association in a case-control study while correcting for population stratification. Genet Epidemiol. 2001;20(1):4–16. Epub 2000/12/19. doi: 10.1002/1098-2272(200101)20:1<4::AID-GEPI2>3.0.CO;2-T. PubMed PMID: 11119293.

4. Pritchard JK, Stephens M, Donnelly P. Inference of population structure using multilocus genotype data. Genetics. 2000;155(2):945–59. Epub 2000/06/03. PubMed PMID: 10835412; PubMed Central PMCID: PMCPMC1461096.

5. Yang H, Wang JR, Didion JP, Buus RJ, Bell TA, Welsh CE, et al. Subspecific origin and haplotype diversity in the laboratory mouse. Nat Genet. 2011;43(7):648–55. Epub 2011/05/31. doi: ng.847 [pii] 10.1038/ng.847. PubMed PMID: 21623374.

6. Price AL, Patterson NJ, Plenge RM, Weinblatt ME, Shadick NA, Reich D. Principal components analysis corrects for stratification in genome-wide association studies. Nat Genet. 2006;38(8):904–9. Epub 2006/07/25. doi: 10.1038/ng1847. PubMed PMID: 16862161.

7. Guenet JL, Bonhomme F. Wild mice: an ever-increasing contribution to a popular mammalian model. Trends Genet. 2003;19(1):24–31. Epub 2002/12/21. doi: S0168952502000070 [pii]. PubMed PMID: 12493245.

8. Reuveni E, Birney E, Gross CT. The consequence of natural selection on genetic variation in the mouse. Genomics. 2010;95(4):196–202. Epub 2010/02/23. doi: S0888-7543(10)00037-6 [pii] 10.1016/j.ygeno.2010.02.004. PubMed PMID: 20171270.

9. Kang HM, Zaitlen NA, Wade CM, Kirby A, Heckerman D, Daly MJ, et al. Efficient control of population structure in model organism association mapping. Genetics. 2008;178(3):1709–23. Epub 2008/04/04. doi: 178/3/1709 [pii] 10.1534/genetics.107.080101. PubMed PMID: 18385116.

10. Sul JH, Martin LS, Eskin E. Population structure in genetic studies: Confounding factors and mixed models. PLoS Genet. 2018;14(12):e1007309. Epub 2018/12/28. doi: 10.1371/journal.pgen.1007309. PubMed PMID: 30589851; PubMed Central PMCID: PMCPMC6307707.

11. Beck JA, Lloyd S, Hafezparast M, Lennon-Pierce M, Eppig JT, Festing MF, et al. Genealogies of mouse inbred strains. Nature Genetics. 2000;24(1):23–5. Epub 1999/12/30. doi: 10.1038/71641. PubMed PMID: 10615122.

12. Zheng M, Dill D, Peltz G. A better prognosis for genetic association studies in mice. Trends Genet. 2012;28(2):62–9. Epub 2011/11/29. doi: 10.1016/j.tig.2011.10.006. PubMed PMID: 22118772; PubMed Central PMCID: PMC3268904.

13. Liao G, Wang J, Guo J, Allard J, Chang J, Nguyen A, et al. In Silico Genetics: Identification of A Novel Functional Element Regulating H2-Ea Gene Expression Science. 2004;306:690–5. PubMed PMID: 3177.

14. Wang J, Peltz G. Haplotype-Based Computational Genetic Analysis in Mice. Computational Genetics and Genomics: New Tools for Understanding Disease. Totowa, New Jersey: Humana Press Inc.; 2005. p. 51–70.

15. Arslan A, Guan Y, Chen X, Donaldson R, Zhu W, Ford M, et al. High Throughput Computational Mouse Genetic Analysis BioRxiv. 2020; https://biorxiv.org/cgi/content/short/2020.09.01.278465v1.

16. Zhang X, Liu HH, Weller P, Zheng M, Tao W, Wang J, et al. In silico and in vitro pharmacogenetics: aldehyde oxidase rapidly metabolizes a p38 kinase inhibitor. Pharmacogenomics J. 2011;11(1):15–24. Epub 2010/02/24. doi: 10.1038/tpj.2010.8. PubMed PMID: 20177421.

17. Ren M, Kazemian M, Zheng M, He J, Li P, Oh J, et al. Transcription factor p73 regulates Th1 differentiation. Nature communications. 2020;11(1):1475. Epub 2020/03/21. doi: 10.1038/s41467-020-15172-5. PubMed PMID: 32193462; PubMed Central PMCID: PMCPMC7081339.

18. Donaldson R, Sun Y, Liang D-Y, Zheng M, Sahbaie P, Dill DL, et al. The multiple PDZ domain protein Mpdz/MUPP1 regulates opioid tolerance and opioid-induced hyperalgesia. BMC Genomics. 2016;17.

19. Zhang H, Zheng M, Wu M, Xu D, Nishimura T, Nishimura Y, et al. A Pharmacogenetic Discovery: Cystamine Protects against Haloperidol-Induced Toxicity and Ischemic Brain Injury. Genetics. 2016;203:599–609.

20. Liang DY, Zheng M, Sun Y, Sahbaie P, Low SA, Peltz G, et al. The Netrin-1 receptor DCC is a regulator of maladaptive responses to chronic morphine administration. BMC Genomics. 2014;15(1):345. doi: 10.1186/1471-2164-15-345. PubMed PMID: 24884839.

21. Zheng M, Zhang H, Dill DL, Clark JD, Tu S, Yablonovitch AL, et al. The Role of Abcb5 Alleles in Susceptibility to Haloperidol-Induced Toxicity in Mice and Humans PLoS Medicine. 2015;12(1):e1001782. Epub Jan 27. doi: 10.1371/journal.pmed.100172.

22. Liu HH, Hu Y, Zheng M, Suhoski MM, Engleman EG, Dill DL, et al. Cd14 SNPs regulate the innate immune response. Mol Immunol. 2012;51(2):112–27. Epub 2012/03/27. doi: S0161-5890(12)00136-8 [pii] 10.1016/j.molimm.2012.02.112. PubMed PMID: 22445606.

23. Sorge RE, Trang T, Dorfman R, Smith SB, Beggs S, Ritchie J, et al. Genetically determined P2X7 receptor pore formation regulates variability in chronic pain sensitivity. Nat Med. 2012;18(4):595–9. Epub 2012/03/27. doi: 10.1038/nm.2710. PubMed PMID: 22447075; PubMed Central PMCID: PMC3350463.

24. Peltz G, Zaas AK, Zheng M, Solis NV, Zhang MX, Liu H-H, et al. Next-Generation Computational Genetic Analysis: Multiple Complement Alleles Control Survival After Candida Albicans Infection Infection and Immunity. 2011;79(11):4472–9.

25. Tregoning JS, Yamaguchi Y, Wang B, Mihm D, Harker JA, Bushell ESC, et al. Genetic Susceptibility to the Delayed Sequelae of RSV Infection is MHC-Dependent, but Modified by Other Genetic Loci. J Immunology. 2010;185(6):5384–91.

26. Hu Y, Liang D, Li X, Liu H-H, Zhang X, Zheng M, et al. The Role of IL-1 in Wound Biology Part II: In vivo and Human Translational Studies. Anesthesia & Analgesia. 2010;111(6):1534–42.

27. Hu Y, Liang D, Li X, Liu H-H, Zhang X, Zheng M, et al. The Role of IL-1 in Wound Biology Part I: Murine in Silico and In vitro Experimental Analysis. Anesthesia & Analgesia. 2010;111(6):1525–33.

28. Liu H-H, Lu P, Guo Y, Farrell E, Zhang X, Zheng M, et al. An Integrative Genomic Analysis Identifies Bhmt2 As A Diet-Dependent Genetic Factor Protecting Against Acetaminophen-Induced Liver Toxicity Genome Research. 2010;20:28–35.

29. Chu LF, Liang D-Y, Li X, Sahbaie P, D’Arcy N, Liao G, et al. From Mouse to Man: The 5-HT3 Receptor Modulates Physical Dependence on Opioid Narcotics. Pharmacogenetics and Genomics. 2009;19:193–205.

30. LaCroix-Fralish ML, Mo G, Smith SB, Sotocinal SG, Ritchie JG, Austin JS, et al. The β3 Subunit of the Na+,K+-ATPase Affects Pain Sensitivity. Pain. 2009;144:294–302.

31. Smith SB, Marker CL, Perry C, Liao G, Sotocinal SG, Austin JS, et al. Quantitative trait locus and computational mapping identifies Kcnj9 (GIRK3) as a candidate gene affecting analgesia from multiple drug classes. Pharmacogenetics and Genomics. 2008;18(3):231–41. Epub 2008/02/28. doi: 10.1097/FPC.0b013e3282f55ab201213011-200803000-00008 [pii]. PubMed PMID: 18300945.

32. Zaas AK, Liao G, Chein J, Usuka J, Weinberg C, Shore D, et al. Plasminogen Alleles Influence Susceptibility to Invasive Aspergillosis. PLoS genetics. 2008;4(6):e1000101. Epub 2008/06/21. doi: 10.1371/journal.pgen.1000101. PubMed PMID: 18566672.

33. Liang D, Liao G, Wang J, Usuka J, Guo YY, Peltz G, et al. A Genetic Analysis of Opioid-Induced Hyperalgesia in Mice Anesthesiology. 2006;104:1054–62. PubMed PMID: 4065.

34. Guo YY, Weller PF, Farrell E, Cheung P, Fitch B, Clark D, et al. In Silico Pharmacogenetics: Warfarin Metabolism. Nature Biotechnology. 2006;24:531–6. PubMed PMID: 4063.

35. Rozzo SJ, Allard J, Choubey D, Vyse T, Izui S, Peltz G, et al. Evidence for an interferon-inducible gene, Ifi202, in the susceptibility to Systemic Lupus. Immunity. 2001;15:435–43. PubMed PMID: 2860.

36. Grupe A, Germer S, Usuka J, Aud D, Belknap JK, Klein RF, et al. In silico mapping of complex disease-related traits in mice. Science. 2001;292(5523):1915–8. PubMed PMID: 2824.

37. Guo YY, Liu P, Zhang X, Weller PMM, Wang J, Liao G, et al. In vitro and In silico Pharmacogenetic Analysis in Mice. Proceedings of the National Academy of Sciences. 2007;104:17735–40. PubMed PMID: 4453.

38. Grubb SC, Bult CJ, Bogue MA. Mouse phenome database. Nucleic Acids Res. 2014;42(Database issue):D825–34. doi: 10.1093/nar/gkt1159. PubMed PMID: 24243846; PubMed Central PMCID: PMC3965087.

39. Patterson N, Price AL, Reich D. Population structure and eigenanalysis. PLoS genet. 2006;2(12):e190.

40. Visscher PM, Wray NR, Zhang Q, Sklar P, McCarthy MI, Brown MA, et al. 10 years of GWAS discovery: biology, function, and translation. The American Journal of Human Genetics. 2017;101(1):5–22.

41. Boyden ED, Dietrich WF. Nalp1b controls mouse macrophage susceptibility to anthrax lethal toxin. Nat Genet. 2006;38(2):240–4. Epub 2006/01/24. doi: ng1724 [pii] 10.1038/ng1724. PubMed PMID: 16429160.

42. Yokoyama T, Silversides DW, Waymire KG, Kwon BS, Takeuchi T, Overbeek PA. Conserved cysteine to serine mutation in tyrosinase is responsible for the classical albino mutation in laboratory mice. Nucleic Acids Res. 1990;18(24):7293–8. Epub 1990/12/25. PubMed PMID: 2124349.

43. Doolittle MH, LeBoeuf RC, Warden CH, Bee LM, Lusis AJ. A polymorphism affecting apolipoprotein A-II translational efficiency determines high density lipoprotein size and composition. J Biol Chem. 1990;265(27):16380–8. PubMed PMID: 2118905.

44. Weng W, Brandenburg NA, Zhong S, Halkias J, Wu L, Jiang XC, et al. ApoA-II maintains HDL levels in part by inhibition of hepatic lipase. Studies In apoA-II and hepatic lipase double knockout mice. J Lipid Res. 1999;40(6):1064–70. PubMed PMID: 10357838.

45. Pittler SJ, Keeler CE, Sidman RL, Baehr W. PCR analysis of DNA from 70-year-old sections of rodless retina demonstrates identity with the mouse rd defect. Proc Natl Acad Sci U S A. 1993;90(20):9616–9. PubMed PMID: 8415750; PubMed Central PMCID: PMCPMC47620.

46. Atwell S, Huang YS, Vilhjalmsson BJ, Willems G, Horton M, Li Y, et al. Genome-wide association study of 107 phenotypes in Arabidopsis thaliana inbred lines. Nature. 2010;465(7298):627–31. Epub 2010/03/26. doi: 10.1038/nature08800. PubMed PMID: 20336072; PubMed Central PMCID: PMCPMC3023908.

47. Ghazalpour A, Rau CD, Farber CR, Bennett BJ, Orozco LD, van Nas A, et al. Hybrid mouse diversity panel: a panel of inbred mouse strains suitable for analysis of complex genetic traits. Mamm Genome. 2012;23(9-10):680–92. Epub 2012/08/16. doi: 10.1007/s00335-012-9411-5. PubMed PMID: 22892838.

48. Chick JM, Munger SC, Simecek P, Huttlin EL, Choi K, Gatti DM, et al. Defining the consequences of genetic variation on a proteome-wide scale. Nature. 2016;534(7608):500–5. doi: 10.1038/nature18270. PubMed PMID: 27309819.

49. Chesler EJ, Miller DR, Branstetter LR, Galloway LD, Jackson BL, Philip VM, et al. The Collaborative Cross at Oak Ridge National Laboratory: developing a powerful resource for systems genetics. Mamm Genome. 2008;19(6):382–9. Epub 2008/08/22. doi: 10.1007/s00335-008-9135-8. PubMed PMID: 18716833.

50. Belknap JK, Crabbe JC. Chromosome mapping of gene loci affecting morphine and amphetamine responses in BXD recombinant inbred mice. Ann N Y Acad Sci. 1992;654:311–23. Epub 1992/06/28. doi: 10.1111/j.1749-6632.1992.tb25977.x. PubMed PMID: 1632590.

51. Zhang X, Liu H-H, Weller P, Tao W, Wang J, Liao G, et al. In Silico and In Vitro Pharmacogenetics: Aldehyde Oxidase Rapidly Metabolizes a p38 Kinase Inhibitor. The Pharmacogenomics Journal. 2011;11(1):15–24. Epub 2010/02/24. doi: tpj20108 [pii] 10.1038/tpj.2010.8.

52. Wainberg M, Sinnott-Armstrong N, Mancuso N, Barbeira AN, Knowles DA, Golan D, et al. Opportunities and challenges for transcriptome-wide association studies. Nat Genet. 2019;51(4):592–9. Epub 2019/03/31. doi: 10.1038/s41588-019-0385-z. PubMed PMID: 30926968; PubMed Central PMCID: PMCPMC6777347.

53. Purcell S, Neale B, Todd-Brown K, Thomas L, Ferreira MA, Bender D, et al. PLINK: a tool set for whole-genome association and population-based linkage analyses. Am J Hum Genet. 2007;81(3):559–75. Epub 2007/08/19. doi: 10.1086/519795. PubMed PMID: 17701901; PubMed Central PMCID: PMCPMC1950838.

54. Hammerschlag AR, de Leeuw CA, Middeldorp CM, Polderman TJC. Synaptic and brain-expressed gene sets relate to the shared genetic risk across five psychiatric disorders. Psychol Med. 2019:1–11. Epub 2019/07/23. doi: 10.1017/S0033291719001776. PubMed PMID: 31328717.

55. de Leeuw CA, Neale BM, Heskes T, Posthuma D. The statistical properties of gene-set analysis. Nat Rev Genet. 2016;17(6):353–64. Epub 2016/04/14. doi: 10.1038/nrg.2016.29. PubMed PMID: 27070863.

56. Watanabe K, Umicevic Mirkov M, de Leeuw CA, van den Heuvel MP, Posthuma D. Genetic mapping of cell type specificity for complex traits. Nature communications. 2019;10(1):3222. Epub 2019/07/22. doi: 10.1038/s41467-019-11181-1. PubMed PMID: 31324783; PubMed Central PMCID: PMCPMC6642112.

57. Yang J, Lee SH, Goddard ME, Visscher PM. GCTA: a tool for genome-wide complex trait analysis. Am J Hum Genet. 2011;88(1):76–82. Epub 2010/12/21. doi: 10.1016/j.ajhg.2010.11.011. PubMed PMID: 21167468; PubMed Central PMCID: PMCPMC3014363.

58. Yang J, Lee SH, Goddard ME, Visscher PM. GCTA: a tool for genome-wide complex trait analysis. The American Journal of Human Genetics. 2011;88(1):76–82.

59. Purcell S, Neale B, Todd-Brown K, Thomas L, Ferreira MA, Bender D, et al. PLINK: a tool set for whole-genome association and population-based linkage analyses. The American Journal of Human Genetics. 2007;81(3):559–75.

60. Benjamini Y, Hochberg Y. Controlling the false discovery rate: a practical and powerful approach to multiple testing. Journal of the royal statistical society Series B (Methodological). 1995;57:289–300.

