## Supplemental Information for "The Effect of Population Structure on Murine Genome-Wide Association Studies"

**Table S1.** The full name of the mouse inbred strain, abbreviation used in the paper, and the Mouse Genome Informatics identification (MGI ID) for the 49 inbred strains used in this study.

| Strain Number | Strain Name | Abbreviation | MGI ID |
|---|---|---|---|
| 1 | 129P2/OlaHsd | 129P2 | 2164147 |
| 2 | 129S1/SvImJ | 129S1 | 3037980 |
| 3 | 129S5/SvEvBrd | 129S5 | 3487126 |
| 4 | A/J | A/J | 2159747 |
| 5 | AKR/J | AKR | 2159745 |
| 6 | B10.D2-H2/oSnJ | B10 | 2160070 |
| 7 | BALB/cJ | BALB | 2159737 |
| 8 | BTBR T<+> Itpr3<tf>/J | BTBR | 2162761 |
| 9 | BUB/BnJ | BUB | 2159907 |
| 10 | C3H/HeJ | C3H | 2159741 |
| 11 | C57BL/6J | C57BL/6J | 3028467 |
| 12 | C57BL/6NJ | C57BL6NJ | 3056279 |
| 13 | C57BL/10J | C57BL10J | 2159754 |
| 14 | C57BR/cdJ | C57BRcd | 2159792 |
| 15 | C57L/J | C57LJ | 2159746 |
| 16 | C58/J | C58 | 2159755 |
| 17 | CAST/EiJ | CAST | 2159793 |
| 18 | CBA/J | CBA | 2159756 |
| 19 | CE/J | CEJ | 2159757 |
| 20 | DBA/1J | DBA1J | 2159759 |
| 21 | DBA/2J | DBA | 2684695 |
| 22 | FVB/NJ | FVB | 2163709 |
| 23 | I/LnJ | ILNJ | 2159844 |
| 24 | KK/HlJ | KK | 2161953 |
| 25 | LG/J | LGJ | 2159748 |
| 26 | LP/J | LPJ | 2159761 |
| 27 | MA/MyJ | MAMy | 2159846 |
| 28 | MOLF/EiJ | MOLF | 2159862 |
| 29 | MRL/MpJ | MRL | 2160037 |
| 30 | NOD/ShiLtJ | NOD | 2162056 |
| 31 | NON/ShiLtJ | NON | 2163530 |
| 32 | NU/J | NUJ | 2161860 |
| 33 | NZB/BlNJ | NZB | 2180844 |
| 34 | NZO/HlLtJ | NZO | 2173835 |
| 35 | NZW/LacJ | NZW | 2159914 |
| 36 | P/J | PJ | 2159762 |
| 37 | PL/J | PLJ | 2159749 |
| 38 | PWD/PhJ | PWD | 2163136 |
| 39 | PWK/PhJ | PWK | 2160654 |
| 40 | RF/J | RFJ | 2159750 |
| 41 | RHJ/LeJ | RHJ | 2162860 |
| 42 | RIIIS/J | RIIIS | 2159809 |
| 43 | SEA/GnJ | SEA | 2159763 |
| 44 | SJL/J | SJL | 2159739 |
| 45 | SM/J | SMJ | 2159787 |
| 46 | SPRET/EiJ | SPRET | 2160671 |
| 47 | ST/bJ | ST | 2159751 |
| 48 | SWR/J | SWR | 2180845 |
| 49 | WSB/EiJ | WSB | 2160667 |

**Table S2.** The significance test results generated by the EIGENSOFT/smartpca program for the first two PCs used for PS characterization are shown. The number of strains, the PCs used (PC1, PC2), the Tracy-Widom (TW)-statistic and the p-value for each PC is shown. The figure that contains the corresponding graph for the analyzed strain groupings is also indicated. A TW statistic with p-value <0.05 indicates the PS is well captured by the PC. The ANOVA analysis for each paired sub-population (determined by IBS matrix) measures the overall genetic differentiation between the two strain groups. (**A, B**) The TW analysis of the groupings of all 49 inbred strains shows that a significant PS is characterized by the first two PCs (**A**); and the inter-group ANOVA test results verify that the sub-populations identified by the PCA are genetically different from each other (**B**). (**C**) The TW results for the 22 most commonly used panels of inbred strains (range: 12 to 33), which were repeatedly used in the indicated number of MPD datasets, reveals that these commonly used strain panels do not have significant PS. The TW p-values decrease as the number of inbred strains in the panel increase, which indicates that the chance of identifying PS increases with the number of strains included in a dataset.

**Table S2A**

| Number of Strains | PC | TW-statistic | p-value | Visualization |
|---|---|---|---|---|
| 49 | 1 | 12.54 | 1.03E-14 | Figure 1 |
| | 2 | 6.04 | 1.75E-6 | |

**Table S2B**

| Population 1 | Population 2 | p-value |
|---|---|---|
| Pop1 | Pop2 | 1.11E-15 |
| Pop1 | Pop3 | 4.22E-26 |
| Pop1 | Pop4 | 7.48E-14 |
| Pop2 | Pop3 | 4.98E-11 |
| Pop2 | Pop4 | 9.35E-10 |
| Pop3 | Pop4 | 1.19E-19 |

**Table S2C**

| Number of Strains | Datasets | PC | TW-statistic | p-value | Visualization |
|---|---|---|---|---|---|
| 12 | 48 | 1<br>2 | -0.77<br>-0.30 | 0.35<br>0.23 | Figure S2A |
| 13 | 17 | 1<br>2 | 0.02<br>-1.94 | 0.16<br>0.71 | |
| 14 | 23 | 1<br>2 | -0.95<br>-0.70 | 0.40<br>0.33 | |
| 15 | 34 | 1<br>2 | 0.13<br>-0.77 | 0.15<br>0.35 | |
| 16 | 118 | 1<br>2 | 0.28<br>-0.64 | 0.12<br>0.31 | |
| 18 | 44 | 1<br>2 | 0.91<br>-1.24 | 0.06<br>0.49 | |
| 19 | 18 | 1<br>2 | -0.91<br>-0.54 | 0.39<br>0.29 | |
| 20 | 94 | 1<br>2 | -0.80<br>-1.50 | 0.36<br>0.57 | Figure S2B |
| 21 | 11 | 1<br>2 | -1.97<br>-1.02 | 0.72<br>0.42 | |
| 22 | 18 | 1<br>2 | -1.37<br>-0.88 | 0.53<br>0.38 | |
| 23 | 44 | 1<br>2 | -0.35<br>-1.94 | 0.24<br>0.71 | Figure S3A |
| 24 | 178 | 1<br>2 | -0.07<br>-0.82 | 0.18<br>0.36 | Figure S4A |
| 25 | 78 | 1<br>2 | 0.08<br>-0.47 | 0.15<br>0.27 | Figure S4B |
| 26 | 30 | 1<br>2 | 0.95<br>-0.84 | 0.05<br>0.37 | |
| 27 | 221 | 1<br>2 | 0.74<br>0.27 | 0.07<br>0.12 | Figure S3B |
| 28 | 100 | 1<br>2 | -0.09<br>-1.12 | 0.18<br>0.45 | Figure S3C |
| 29 | 58 | 1<br>2 | 1.52<br>0.18 | **0.02**<br>0.14 | Figure S4D |
| 30 | 34 | 1<br>2 | 1.42<br>0.38 | **0.03**<br>0.11 | Figure S4E |
| 31 | 55 | 1<br>2 | 0.97<br>-0.42 | 0.05<br>0.26 | |
| 32 | 50 | 1<br>2 | 1.42<br>0.17 | **0.03**<br>0.14 | Figure S4F |
| 33 | 61 | 1<br>2 | 0.52<br>0.18 | 0.09<br>0.14 | |

**Table S3.** Summary table showing the number of MPD datasets that measure responses using the indicated number of inbred strains. For our analyses, we selected 2435 MPD datasets that measured a response in 10 or more inbred strains. Of note, 43% of these datasets analyzed 27, 24, 23, or 20 inbred strains.

| Number of Strains | Number of Datasets |
|---|---|
| 10 | 9 |
| 12 | 68 |
| 13 | 48 |
| 14 | 33 |
| 15 | 82 |
| 16 | 136 |
| 18 | 103 |
| 19 | 47 |
| 20 | 159 |
| 21 | 41 |
| 22 | 53 |
| 23 | 162 |
| 24 | 265 |
| 25 | 132 |
| 26 | 88 |
| 27 | 416 |
| 28 | 147 |
| 29 | 138 |
| 30 | 73 |
| 31 | 94 |
| 32 | 80 |
| 33 | 61 |

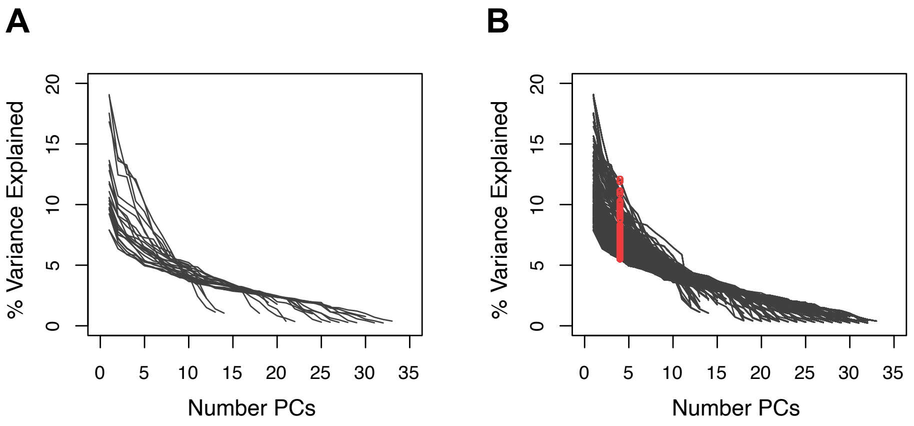


**Figure S1.** The percentage of variance that is explained when the indicated number of principal components (PC) are used to analyze 22 selected MPD datasets (**A**) or all 2435 MPD datasets (with $\geq$10 inbred strains) (**B**). Every MPD dataset analyzed a response in a panel of inbred strains (range 10 to 33 strains). Each individual curve represents a PCA for one mouse population for a dataset. The red circles show the percentage of the variance that is explained by the 4th PC, which ranges from 5.5%-12.1%. The total variance explained ranges between 26% and 59% when the first four PCs are used.

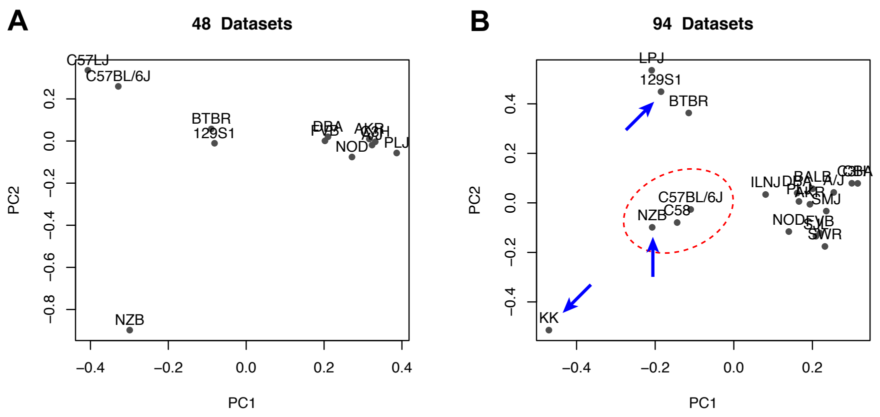


**Figure S2.** PCA plots (using the first two PCs) examine the genetic relationships of the 12 strains that are evaluated in 48 MPD datasets (A) or the 20 inbred strains evaluated in 94 MPD datasets (**B**). (**A**) In this graph, although NZB is separated from the other strains; a single strain cannot by itself be classified as a sub-population. Also, while there appear to be 4 sub-populations in this plot, three of the groups contain only 1 or two strains. (**B**) This graph appears to have 4 sub-groups of strains. However, KK (a global group 2 strain, blue arrow at bottom) forms a strain group that is separated from other group 2 strains (NZB, BTBR, 129S1). Moreover, global group 2 (NZB) and group 1 (C57BL/6J, C58) strains form one group in this graph (red circle). Overall, when a small number of strains are evaluated, the sub-population structure is highly variable, and it can be significantly altered by adding or deleting a single strain.

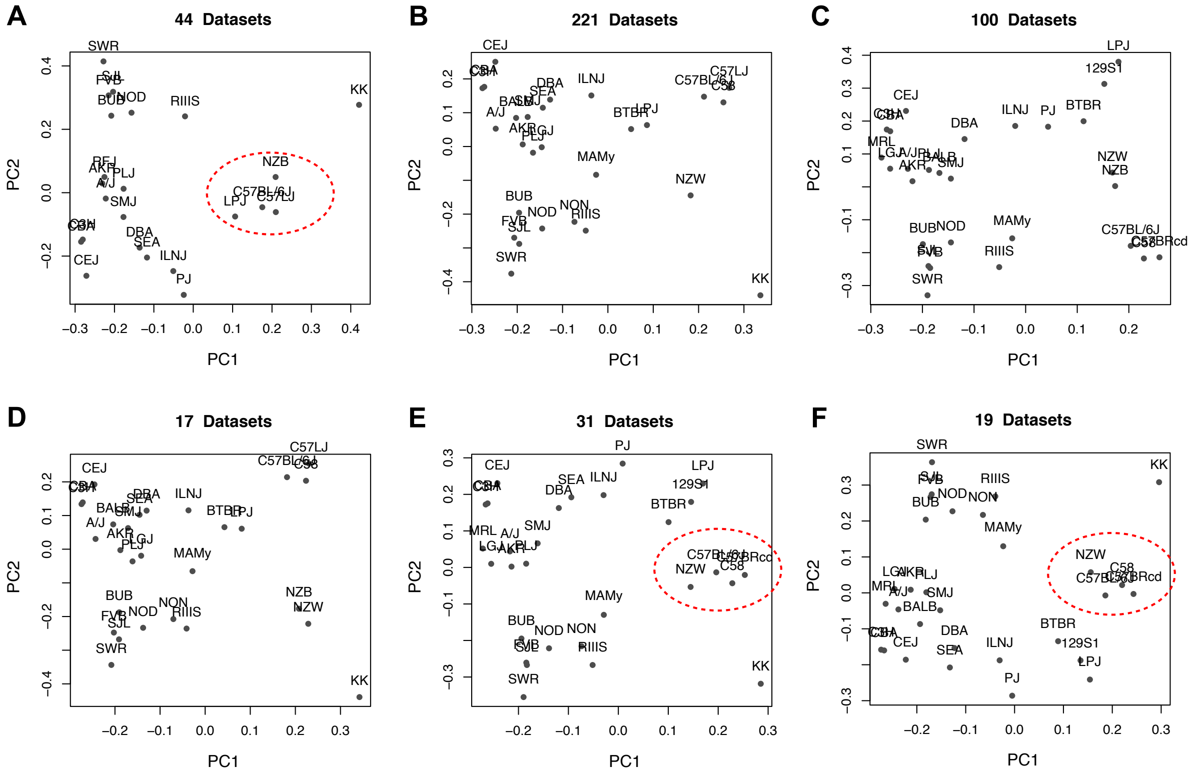


**Figure S3.** Six commonly used panels of inbred strains, whose responses were measured in 432 MPD datasets, lack population sub-structure. Scatter plots, which show the first two PCs, were generated to examine population structure among these commonly used strains. These MPD datasets measure responses (from A to F) in the same set of 23 strains (44 datasets), 27 strains (221 datasets); 28 strains (100 datasets) and a second set of 17 datasets used a different 28 strain panel; 29 strains (31 datasets); and 30 strains (19 datasets). The red circles indicate groupings where the local population sub-structure violates the global PS: group 1 (C57BL related) strains are intermixed group 2 (NZB, NZW, LPJ) strains.

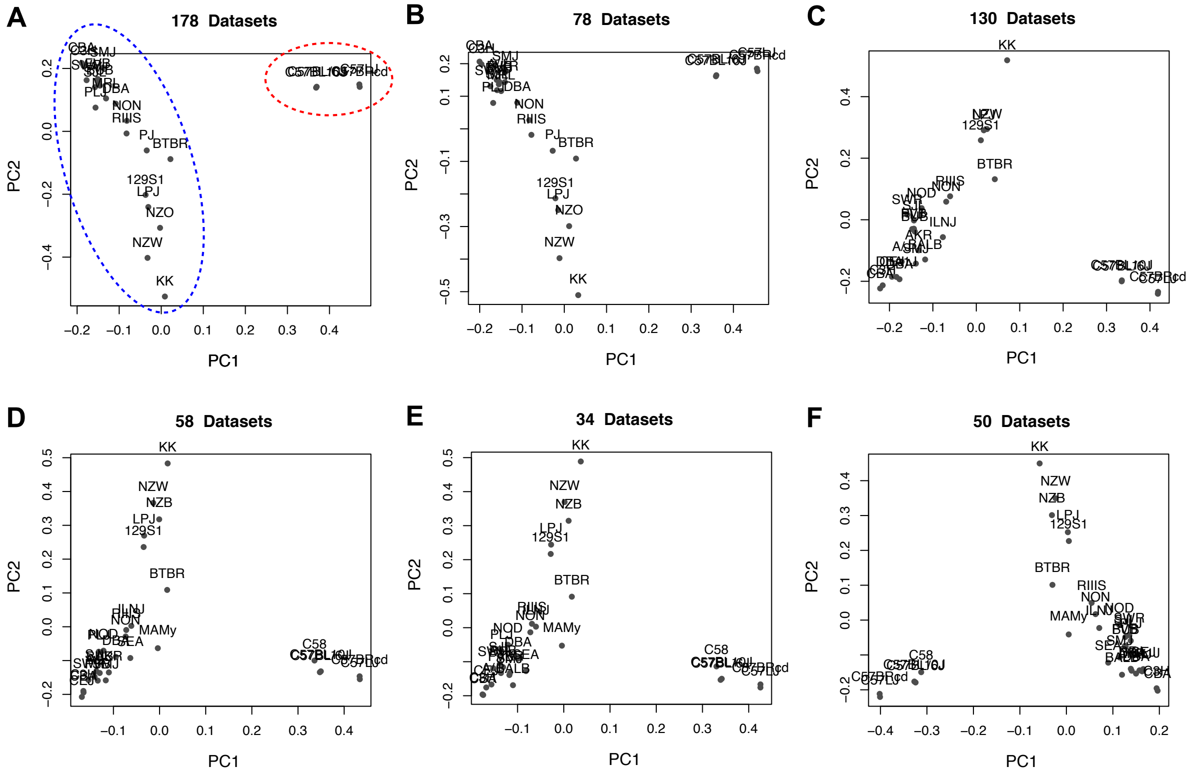


**Figure S4.** Six commonly used panels of inbred strains, which were used to generate 528 MPD datasets, have some degree of PS. Scatter plots showing the first two PCs were generated to examine population structure among the 24 to 32 inbred strains evaluated in these 535 MPD datasets. Responses in these datasets were measured (from A to F) using the same set of 24 strains (178 datasets); 25 strains (78 datasets); 27 strains (130 datasets), 29 strains (58 datasets); 30 strains (34 datasets); and 32 strains (50 datasets). The group 1 strains (C57BL related) are clearly separated from other strains (red circle, panel A). However, the global group 2 and group 3 strains are broadly distributed in these graphs, and they cannot be separated into distinct sub-groups (see blue circle).
